# Cdc23/Mcm10 Primase Generates the Lagging Strand-Specific Ribonucleotide Imprint in Fission Yeast

**DOI:** 10.1101/303180

**Authors:** Balveer Singh, Kamlesh K Bisht, Udita Upadhyay, Avinash Chandra Kushwaha, Jagpreet Singh Nanda, Suchita Srivastava, Amar J.S. Klar, Jagmohan Singh

## Abstract

The developmental asymmetry of fission yeast daughter cells derives from inheriting “older Watson” versus “older Crick” DNA strand from the parental cell, strands that are complementary but not identical with each other. A novel DNA strand-specific “imprint”, installed during DNA replication at the mating-type locus (*mat1*), imparts competence for cell type inter-conversion to one of the two chromosome replicas. The biochemical nature of the imprint and the mechanism of its installation are still not understood. The catalytic subunit of DNA Polymerase α (Polα) has been implicated in the imprinting process. Based on its known biochemical function, Polα might install the *mat1* imprint during lagging strand synthesis. The nature of the imprint is not clear: it is either a nick or a ribonucleotide insertion. Our investigations do not support a role of Polα in nicking through putative endonuclease domains but confirm its role in installing an alkali-labile moiety as the imprint. A detailed genetic and molecular analysis reveals a direct role of the Cdc23/Mcm10 primase activity in installing the imprint in cooperation with Polα and Swi1.

## INTRODUCTION

In *Schizosaccharomyces pombe*, the mating-type region comprises three loci located on chromosome II: *mat1M* or *P*, *mat2P* and *mat3M* (Fig. 1A). The *mat1* cassette is expressed and it dictates the Plus or Minus sex/cell type to the cell. The *mat2P* and *mat3M* cassettes, which are transcriptionally silenced by an epigenetic mechanism, function as master copies for switching *mat1*. Switching occurs by highly regulated recombination through transposition/substitution of the *mat1* allele with the opposite mating-type information, copied from either *mat2P* or *mat3M* cassettes [1-4]. The mating-type switching process is exquisitely regulated in very interesting ways. First, of the four “granddaughter” cells derived from a single cell, only one cell switches in nearly 90% of pedigrees [2,5,6]. This pattern results from asymmetric cell division occurring in each of the two consecutive generations in the progeny of a single cell. The switching program is initiated by a novel “imprint,” that is installed in specific DNA strand during replication of the *mat1* locus on chromosome II [1,2]. In the following cell division, the imprint consummates into a switch but only in one of the sister chromatids during *mat1* replication. The *mat1*-switching event removes the imprint. The *mat1* locus in the chromosome is replicated only in one direction. Chromosomal inversion of the *mat1* “cassette” abolishes imprinting, which is partially restored by genetic manipulations promoting *mat1* replication in the opposite direction [7]. Thus, replication of a specific *mat1* strand specifically by the lagging-strand replication complex is critical for generating the imprint. The imprinting process requires three genes [8]: *swi1* [9], *swi3* [10] and *swi7/pol*α [11]. Mechanistically, Swi1 and Swi3 create a replication pause at the imprint site. They also block replication forks originating from the centromeric side of *mat1* [9]. The single *swi7-1/pol*α imprinting-deficient mutant, however, shows a normal pause and normal replication fork block, and is, therefore, defective at some other undefined step in imprinting pathway [9].

**Figure 1.**
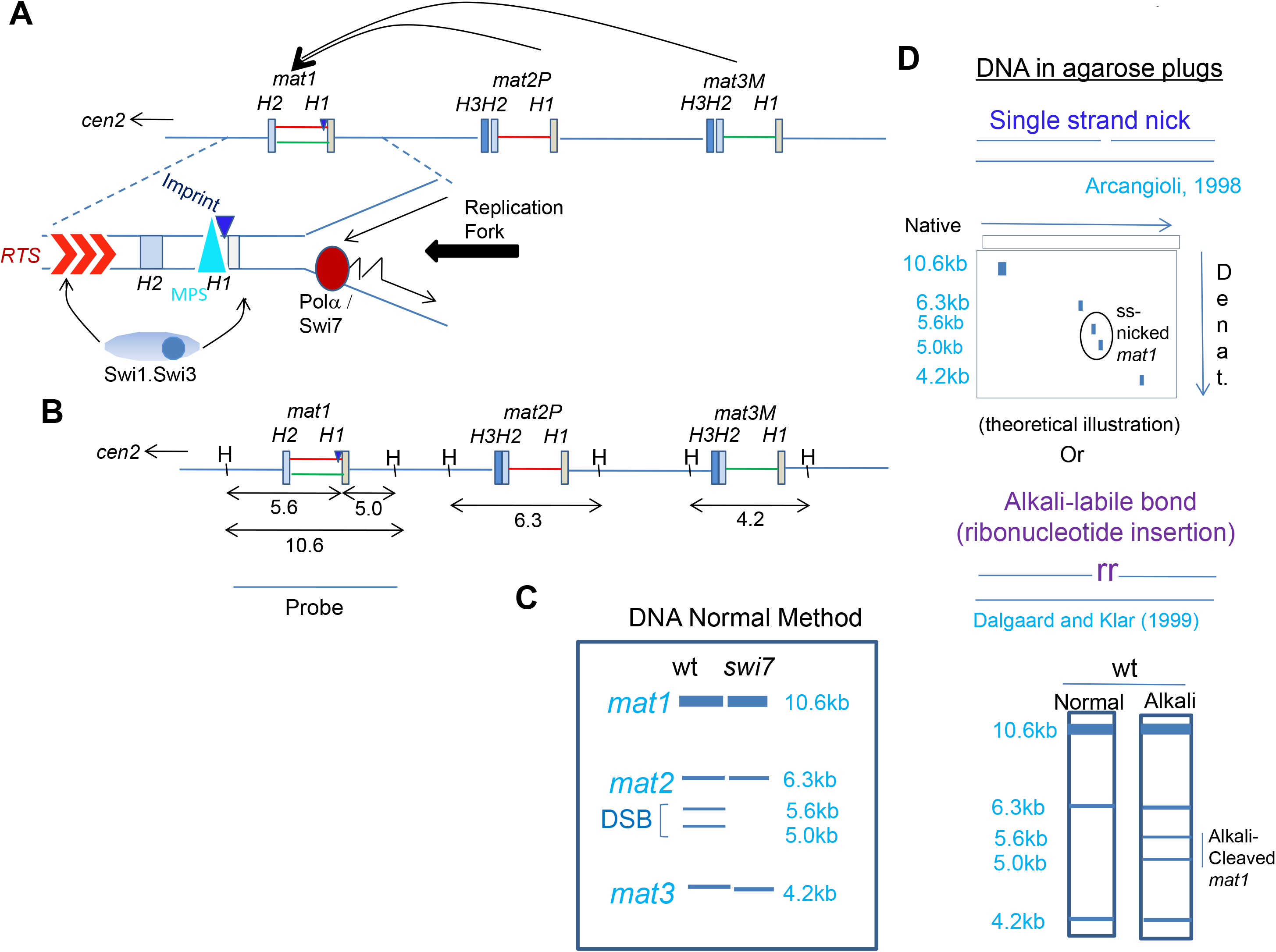
Schematic diagram depicting the organization of mating-type loci in *S. pombe*. **A**, The loci *mat1, mat2* and *mat3* are located in ~30kb region of chromosome 2. They comprise short conserved homology regions H1 to H3, which flank the ~1.1 kb allele-specific sequences. Following imprinting occurring at the boundary of *mat1* and the allele-specific region (dark blue triangle), a copy of the donor locus (*mat2* or *mat3*) is transposed to the *mat1* locus resulting in its switching by gene replacement. Replication fork progression from the centromere distal direction is met by a pause site (MPS1, blue triangle; 9), while fork from the left side encounters the replication termination sequence (RTS; 36). **B**, A schematic showing the *Hin*dIII restriction pattern of the mating type region with the corresponding picture observed following Southern blot hybridization using the *mat1P* or *M*fragment of 10.6 as a probe (C), wherein *mat1, mat2* and *mat3* loci migrate at the positions of 10.6, 6.2 and 4.3 kb, respectively. Occurrence of imprint at *mat1* generates a fragile site, which appears as a double strand break (DSB) generating the bands of 5.6 and 5.0 kb when DNA is prepared by the conventional method (left lane). Due to lack of imprint no DSB is seen in *swi1, swi3* or *swi7* mutants. **D**, Schematic representation of the methodology used to detect a nick as the imprint, which can be visualized by 2-dimensional gel electrophoresis. (Top panel) DNA is prepared in plugs and then resolved by acrylamide gel electrophoresis in the first dimension, while the 2^nd^ dimension is run in a denaturing gel. Alternatively, samples embedded in agarose are digested with *Hin*dIII and then subjected to electrophoresis in native agarose gel without or with alkali-treatment (lower panel).

However, the nature of the imprint remains unresolved; it is thought to be either a site- and strand-specific nick [12] or an alkali-labile, RNase-sensitive modification, consisting of one or two ribonucleotides incorporated in *mat1* DNA [7,13,14]. The imprint creates a DNA fragile site, which is artifactually converted into double-strand break (DSB) due to hydrodynamic shear when extracting DNA from cells [7,12]. Therefore, the imprint level is usually determined by quantifying the DSB at *mat1* through Southern blot analysis.

The *swi7* gene, encoding the catalytic subunit of Polα, is inherently required much more often in the lagging-than the leading-strand DNA synthesis during chromosome replication [11]. The biochemical role of Polα/Swi7 in generating the imprint has remained elusive. This is because it encodes a gene which is essential for viability, thus limiting its analysis; only one allele, *swi7-1/Pol*α, mapping to the carboxy-terminal region (G1116E) of the catalytic subunit of Polα, is known to affect imprinting [11]. Notably, the imprinting event occurs only on the newly synthesized lagging strand during S phase [15]. Since the Polα/Primase complex can synthesize and extend an RNA moiety on DNA template, Polα/Swi7 is a plausible candidate for installing the imprint in the form of ribonucleotide(s) insertion at *mat1*, through the primase subunit. Alternatively, Polα-catalyzed DNA nicking may constitute the imprint [12].

## RESULTS AND DISCUSSION

### Polα catalytic subunit does not play a direct role in Imprinting

This work was initiated to determine whether the catalytic subunit of Polα may be directly involved in generating the imprint in the form of a nick at the *mat1* locus. It was shown earlier that the DNA prepared by the normal method causes hydrodynamic shearing resulting in conversion of a nick into double strand break at the imprint site [12]. As a result the *mat1 Hin*dIII fragment of 10.6kb is split into two bands of 5.6 and 5.0kb (Fig. 1B, 1C). According to this study, preparation of the DNA in agarose plugs avoids the hydrodynamic shear and *mat1* DNA shows a nick in the top DNA strand as the putative imprint [12]. Assuming that Polα may act as an endonuclease, we analyzed the sequence of Polα for the presence of endonuclease motifs and found motifs similar to the putative intein endonuclease [16-18] and restriction endonucleases [19] (Expanded View, Fig. EV1A-D). We investigated whether these motifs play a role in imprinting. However, we found that the plasmids containing the mutated *Pol*α gene complemented the *swi7-1* mutation, ruling out a role of Pola as an endonuclease (Expanded View, Fig. EV1A-D). Surprisingly, we found that a catalytically dead version of Polα (D984N mutant), which is unable to extend the RNA primers synthesized by the Polα-primase subunit [20], was actually proficient in restoring the imprint (detected as DSB using DNA prepared by the normal method) when transformed into *swi7-1* mutant. The mutant plasmid did not complement the imprinting defect in the *swi1* and *swi3* mutants [Expanded View; Fig. EV2A].

As we were unable to identify any endonuclease function of Polα in explaining the DNA nick, we considered the alternative possibility that the imprint may be due to the presence of ribonucleotide/s in the DNA chain, which is detectable as an alkali-labile bond [7]. Indeed, a break was indicated by a broad band at the putative imprint site, at 5.0-5.6kb region, in Southern blots of alkali-treated plugs of DNA digested with *Hin*dIII [Expanded View; Fig. EV2B, right panel]. Thus, the imprinting step does involve a ribonucleotide insertion as a step distinct from the replication function of Polα.

These results created a conundrum for us: how might the catalytically dead Polα complement the imprinting defect of *swi7-1* mutant? The *Polα^D984N^* mutation in the evolutionarily conserved Asp residue does not alter either the stability or assembly of the mutant Polα-primase complex [20-21]. Notably, the catalytic subunit mutant protein is unable to further extend RNA primers synthesized by the primase subunit [20-21]. We envisaged two possibilities: either the Polα^Swi7-1^ mutant protein complex is defective in the primase activity or in utilizing the RNA primer synthesized by the RNA primase subunit for DNA synthesis. To distinguish between these possibilities, we tested the imprinting and switching efficiency of temperature sensitive mutants in primase subunits of Polα: those of *spp1-4, spp1-14* alleles of *spp1* and *spp2-8* allele of *spp2* genes (Fig. EV3A) [22-24]. Although, somewhat reduced efficiency of switching was observed, especially in spp1-4 mutant (Fig. EV3B), the level of imprint/DSB was affected in these mutants to an extent similar to the wt strain when cultured at semi-permissive growth temperature of 34°C (Fig. EV3C). Thus, these data do not support a role of the Polα-primase subunit in *mat1* imprinting. However, the extremely low rate of switching in the *spp1-4* mutant at 34°C may be due to lower efficiency of utilization of the imprint for switching, as observed earlier in case of *swi2, swi5* and *swi6* mutants [8].

### Other Mcm components do not play role in imprinting

Our findings indicated a role of DNA replication initiation in imprinting. Next, we queried whether the components of MCM-helicase complex, essential for DNA replication initiation, are required for imprinting (Fig. EV4A. We observed normal iodine staining of colonies of mutants in *mcm2* (*cdc19-p1*) [25], *mcm4* (*cdc21-M68*) [26], *mcm5* (*nda4-108*) [27] and *mcm6* (*mis5-268*) [28], in homothallic (*h^90^*) background indicative of normal switching at permissive temperature (25°C) and semi-permissive temperature (30°C) (Expanded View; Fig. EV4B). This result argues against the role of earlier steps of replication initiation in imprinting.

### The Cdc23/SpMcm10 primase performs previously unknown imprinting function at ***mat1***

Because of a reported primase-like function of Cdc23/SpMcm10, we next investigated the efficiency of switching and *mat1* imprinting of *cdc23* mutants (Fig. 2A). Notably, the *cdc23-M30* and *cdc23-M36* mutations are located in its Mcm/Polα-interacting domain [29,30]. Both *cdc23-M30* and *cdc23-M36* mutants showed normal switching (Fig. 2B, top panel) and DSB/imprint (Fig. 2D, left panel) in cells grown at permissive temperature (25°C) but showed reduced switching and DSB at 30°C semipermissive growth temperature (Fig. 2B, lower panel; Fig 2D, right panel, 2E). In contrast, the *cdc23-1E2* mutant, its mutation also mapping to the Mcm- and Polα-interacting domain (Fig. 2A), showed low switching efficiency (Fig. 2B. lower panel) and much reduced DSB/imprint at both growth temperatures (Fig. 2D, 2E). Importantly, *cdc23-M36* mutant showed an alkali-labile bond at *mat1* at growth temperature of 25°C but not at 30°C semi-permissive temperature (Fig. 2C, 2F, right panel; compare lanes 6 and 8; 2G). Thus, Mcm- and Polα-interacting domain of Cdc23 plays a role in installing the alkali-labile imprint.

**Figure 2.**
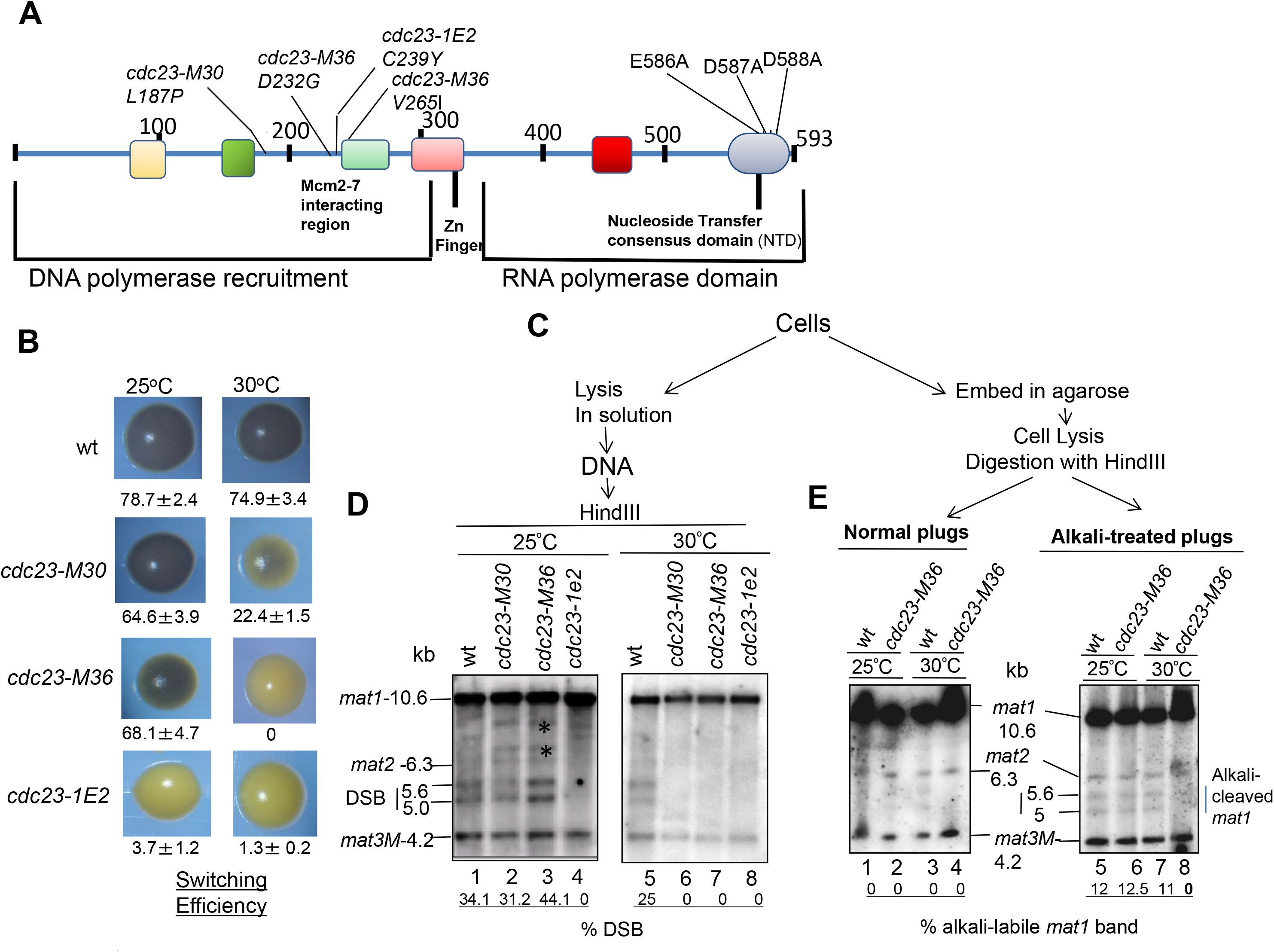
Role of Cdc23/SpMcm10 in Imprinting at the *mat1* locus. **A**, Structure of Cdc23 protein, indicating the N-terminal domain required for binding with ssDNA and DNA Polα, the Zinc finger domain and the C-terminal domain performing the RNA polymerase function. Locations of mutations M30, M36 and 1E2 in the N-terminal domain and D588A in the C-terminal RNA polymerase domain are shown (adapted from Ref. 29). **B**, Iodine-staining phenotypes of the indicated strains in *h^90^* background after growth on PMA^+^ plates at 25°C or 30°C. **C**, Schematic showing the process of preparation of DNA by normal method or in plugs.* represents bands due to mating type rearrangements. **D**, *cdc23* mutants exhibit imprinting defect. DNA was prepared and analyzed by the standard method. **E**, DNA was prepared in plugs from indicated cultures grown at 25°C or 30°C, without (left panel) and with alkali treatment (right panel) followed by Southern blotting.

Previous structure-function analysis of Cdc23 has revealed three functional domains: Polα-interaction domain, zinc domain and putative primase domain, having similarity to the bacteriophage T7 gene 4 primase (Fig. 2A) [29]. Because the primase mutants are inviable [29], we explored the role of Cdc23 domains in imprinting by genetic complementation experiments. We tested the ability of the high copy number plasmids bearing different *cdc23* mutant alleles to suppress the imprinting defect of the *cdc23-M36* allele. While plasmids bearing wild-type *cdc23^+^*, *cdc23-M30, cdc23-M36, cdc23-1E2* or *cdc23-D587A* complemented switching defect of the *cdc23-M36* mutant grown at 30°C, the primase defective mutant *cdc23-D588A* [29] and *Polα* genes did not (Fig. 3A). Surprisingly, a primase proficient mutant E586A also failed to complement the switching defect (Fig. 3A). Analysis of DNA prepared by conventional method paralleled the complementation data (Fig. 3B), showing that transformants containing wild-type *cdc23^+^*, *cdc23-M30, cdc23-M36, cdc23-1E2* or *cdc23-D587A* complemented the imprinting defect, but *cdc23-D586A*, cdc23-D588A and *Polα* did not (Fig. 3B). Herein, the inability of primase proficient mutant E588A to complement the imprinting defect paralleled its inability to complement the switching defect. This surprising finding was also reported earlier [29] wherein the E588A mutant was found to ineffective in complementing the growth defect of the *mcm10Δ* mutant of *S. cerevisiae*. This discrepancy was ascribed to the effect of E588A mutant on function of the adjoining catalytic site of primase or on protein-protein interactions [29].

**Figure 3.**
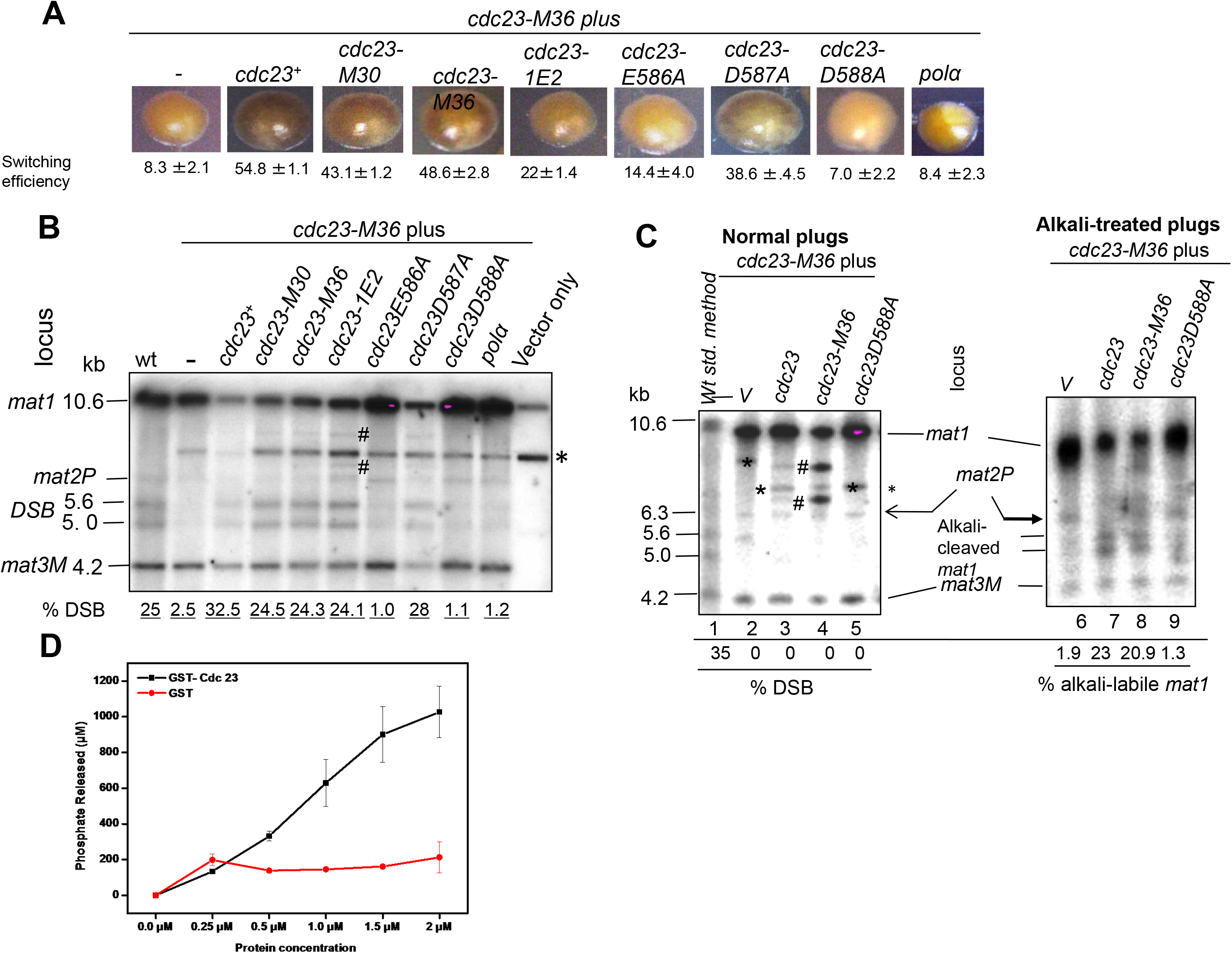
Primase domain of Cdc23 is essential for imprinting. **A**, Iodine-staining phenotypes of *h^90^, cdc23-M36* mutant transformed with plasmids bearing indicated *cdc23* gene mutations and grown on selective medium plates at 30°C. **B-C**, *cdc23* genes having mutation in primase domain fail to complement the imprinting defect of the *cdc23-M36* mutant. DNA was prepared from transformants of *cdc23-M36* mutant with the indicated high copy plasmids containing the indicated genes and grown at 30oC. **B**, DNA was prepared by the conventional method. **C**, DNA was prepared in agarose plugs. Lane labeled wt indicates DNA prepared from wild type strain. Vector only lane shows cross-reacting vectors backbone bands (marked by *) present in all lanes, while bands marked (#) correspond to mating-type rearrangements. DNA was digested with *Hin*dIII and subjected to agarose gel electrophoresis. **C**, DNA prepared in plugs was gel was blotted run without (left panel) and with alkali treatment (right panel) from strains grown at 30°C. After Southern blotting membranes were subjected to hybridization. **D**, Cdc23 possesses primase activity. The amount of PPi released by primase assay was assayed according to [49].

The results with DNA prepared in plugs with and without alkali treatment showed that plasmids bearing *cdc23^+^* and *cdc23-M36* restored the alkali-labile bond at *mat1* in *cdc23-M36* mutants when cultured at 30°C (Fig. 3C, compare lanes 7 and 8 with lanes 3 and 4), while plasmids bearing the primase defective *cdc23-D588A* gene did not (Fig. 3C, lane 9).

As the primase function of Mcm10 has not been demonstrated in other orthologs [31], we tested for the primase activity of the purified recombinant GST-tagged Cdc23. Results showed that GST-Cdc23 does possess primase activity when assayed by measurement of the amount of pyrophosphate released due to the primase activity(Fig. 3D). Taken together, these results confirmed the primase function of Cdc23 from *S. pombe* and identified a novel function of Cdc23 primase in imprinting, a result consistent with the possibility of ribonucleotide(s) constituting the imprint moiety.

### Genetic and biochemical Interactions of Cdc23 with Swi1, Swi3 and Swi7

Polα physically interacts with Cdc23 [30-32]. Since both *cdc23* and *swi7-1/Polα* mutants are defective in imprinting, we tested genetic interaction between their mutations. The double mutant showed much reduced sporulation efficiency, indicating reduced rate of switching on minimal medium at 30°C (Fig. EV5A). This result indicated that both these factors are required at different steps in imprinting. Surprisingly, unlike the single mutants, the double mutant failed to grow on rich media at 30°C (Fig. EV5C).

We also investigated genetic interactions of *cdc23* mutant with *swi1* and *swi3* mutants in switching and viability. Interestingly, *cdc23-M36* mutant showed cumulative effect on switching efficiency in combination with *swi1* and *swi3* mutants on minimal medium at 30°C (Fig. EV5B), suggesting that Cdc23 and Swi1/Swi3 may act at different steps of imprinting. The double mutants of *cdc23* with *swi1* and *swi3* mutants also showed synthetic lethality on rich medium at 30°C (Fig. EV5D). The discrepancy between growth on minimal medium and lack of growth on rich medium at 30oC is surprising. It may reflect that the double mutants affect some important physiological function during vegetative growth but not during starvation.

We further investigated direct biochemical interactions *in vivo* and *in vitro*. Polα^+^p could be readily co-immunoprecipitated with Cdc23^+^-HA but in comparison the mutant Polα^7-1^p was less efficiently co-immunoprecipitated (Fig. 4A). *In vitro* pull-down experiments showed that MBP-Polα^+^ interacted more strongly with GST-Cdc23 than with GST-Cdc23^M36^ (Fig. 4B, panels I and II; Fig. 4C). Likewise, Polα^7-1^ interacted more strongly with Cdc23^+^ than with Cdc23^M36^ (Fig. 4B, panels III and IV; Fig. 4C). Furthermore, Cdc23 interacted with Polα^+^ more strongly than with Polα^7-1^ (Fig. 5B, compare panels I and III, panels II and IV; Fig. 5C). The order of strength of interaction was Polα^+-^ Cdc23^+^ > Polα^Swi7-1^-Cdc23^+^ > Polα^+-^ Cdc23-M36 = Polα^Swi7_1^-Cdc23-M36. As Cdc23 recruits Polα to chromatin [33-35], our results suggest that the recruitment of Polα may be reduced in *swi7-1* and *cdc23-M36* mutants.

**Figure 4.**
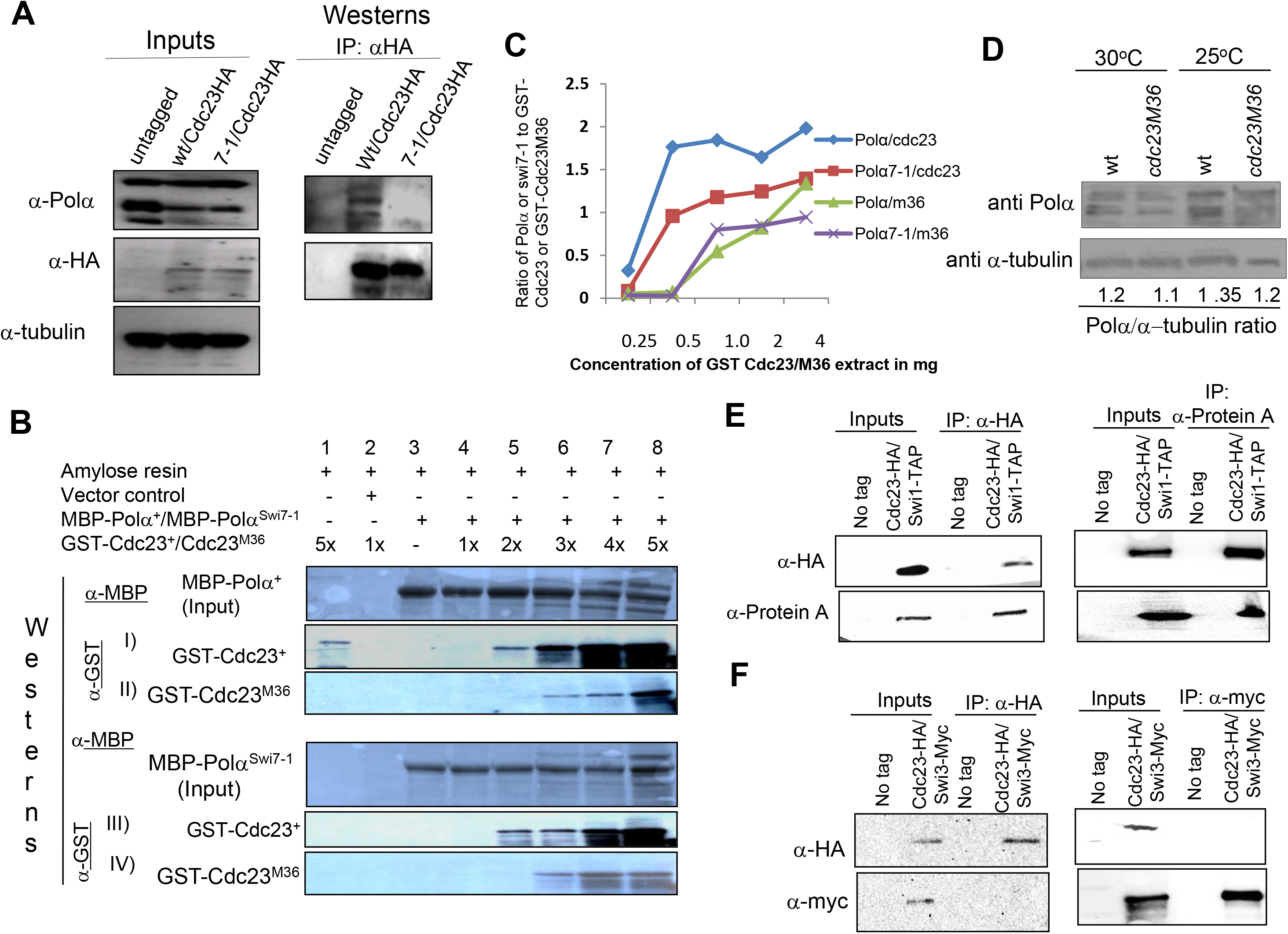
Reduced interaction of Cdc23 and Polα mutant proteins. **A**, Co-IP of HA-tagged Cdc23 with Polα and Polα^swi7-1^. Inputs blot shows equivalent amounts of HA-taged Cdc23, Polα and Polα^swi7-1^ was present in the indicated strains. Western; IP was performed with anti-HA antibody followed by immunoblotting with anti-HA and anti-Polα antibodies. **B,C**, *In vitro* interactions. **B**, Identical amounts of MBP-Polα (panels I and II) and MBP-Polα^swi7-1^ (panels III and IV) were bound to amylose beads and incubated with increasing concentrations of the normalized amounts of GST-Cdc23 (panels I and III) and GST-Cdc23^M36^ (panels II and IV), as indicated. Following SDS-PAGE of the bound proteins, the blots were probed with anti-MBP (panels I and II) and anti-GST antibodies (panels III and IV). **C**, Quantitation of MBP-Polα and MPB-Polα^swi7-1^ binding with Cdc23 and Cdc23^M36^. X axis shows the concentration of Cdc23 or Cdc23^M36^ and the Y axis shows the ratio of Polα or Polα^swi7-1^ to input GST-Cdc23 or GST-Cdc23^M36^. **D**, No deleterious effect of *cdc23M36* mutation on the level of DNA Polα. Equal amounts of proteins prepared from wt and *cdc23-M36* mutant grown at 25°C and 30°C were subjected to SDS-PAGE and western blotting with anti-Polα and anti-a-tubulin antibodies. **E-F**, Coimmunoprecipitation experiment showing that Cdc23 interacts with Swi1 (**E**) but not with Swi3 (**F**) *in vivo*. Strains without tag or the indicated tags for Cdc23 (HA) and Swi1 (TAP) and Swi3 (myc) were used for immunoprecipitation followed by immunoblotting with the indicated antibodies.

**Figure 5.**
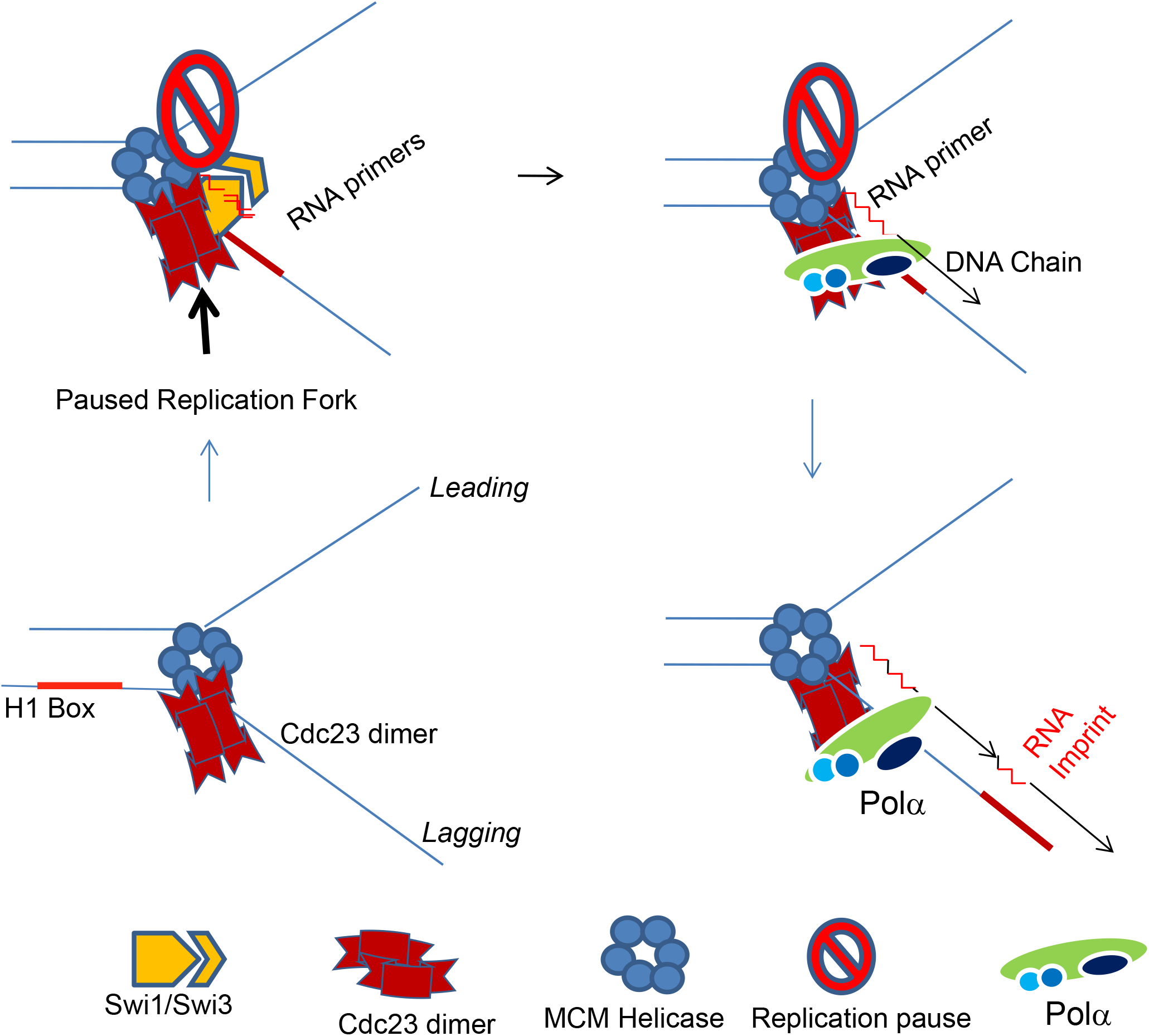
A model visualizing the role of Cdc23 Primase and DNA Polα during lagging strand synthesis in generating the imprint at mat1 locus. Pausing at the imprint site by Swi1 and Swi3 may lead to allow Cdc23 primase to interact with Swi1 and to linger at the pause site for prolonged period during elongation phase of DNA replication. This may allow synthesis a short primer of 2 ribonucleotides length, which is subsequently extended by the catalytic subunit of Polα.

A trivial possibility to explain our genetic results could be reduction in the level of Polα in *cdc23* mutants, as inactivation of Cdc23 homolog, Mcm10, is known to cause rapid loss of Polα in *S. cerevisiae* [33]. However, western blot analysis shows that, compared to the wild type strain, the level of Polα is not reduced significantly in *cdc23-M36* mutant grown at 25°C and 30°C (Fig. 4D).

Interestingly, co-immunoprecipitation experiments also showed that Cdc23 interacts with Swi1 but not Swi3 *in vivo* (Fig. 4E, 4F). Thus, Cdc23 may also act at a common step involving Swi1 and Polα.

Our study was initiated to address the mechanism of action of catalytic subunit of DNA Polα in strand-specific imprinting at the *mat1* locus in *S. pombe*, which imparts competence to switch cell type in the following cell division [1,2]. Swi1 and Swi3 function to produce replication pause at the imprint site [7,9] and at a centromere-proximal replication termination sequence [36]. While Swi1 and Swi3 play an important but indirect role by providing a paused replication fork to promote imprinting, the enzymatic activity catalyzing the imprint was not known. Given its role in lagging strand synthesis [11,37], we investigated whether the catalytic subunit of Polα plays a direct role in imprinting.

As the imprint is either a nick [12] or 1/2 ribonucleotide insertion [7], our initial investigation addressed the possibility of Polα causing a nick. We discounted this possibility because mutations in two potential endonuclease sequence motifs complemented the imprinting defect of the *swi7-1* mutant. Most surprisingly, even a catalytically dead *Polα^D984N^* mutant gene was complementation proficient. Although the Polα^D984N^ mutant protein forms a stable complex with the primase Spp1 and Spp2 subunits, the catalytic subunit fails to extend the RNA primer synthesized by the primase subunit [21,22]. The puzzling effect of the catalytically dead Polα in restoring the imprint motivated us to investigate whether a step of DNA replication, earlier than the Okazaki fragment synthesis, is critical for imprinting. None of the components of the MCM helicase complex was found to involved in imprinting. Likewise, the *Spp1* and *Spp2* primase subunit mutants showed normal imprinting. Finally, mutants of the non-canonical primase *cdc23*/Sp*mcm10* gene were found to be defective in imprinting.

Mcm10/Cdc23 has been studied both in *S. cerevisiae* and *S. pombe. In vitro* studies have shown that the SpCdc23 makes 2-20nt long RNA primers, which are then transferred to and extended by the catalytic subunit of Polα [29]. Notably, it is the N-terminal domain of Cdc23 that recruits Polα to replication fork [29,30]. Taken together with the *in vitro* and *in vivo* studies showing a reduced ability of mutant Cdc23-M36 to interact with Polα, as well as, that of Cdc23 with Polα^7-1^, these results suggest that both interaction of Cdc23 with Polα along with its own primase-like function are required for imprinting (Fig. 5). It is pertinent that even a short, 2-ribonucleotide primer synthesized by Cdc23 can be extended into a DNA chain by Polα [29], As the imprint is constituted by a diribonucleotide insertion [14], we suggest that primers as short as 2 ribonucleotides, that are synthesized by primase domain of Cdc23 at the *mat1* locus, can be extended into a DNA chain by the catalytic domain of Polα; Polα, in turn, is recruited through N-terminal domain of Cdc23 [29,30]. It is puzzling how such an event could occur at the pause site. In this regard it has been shown that multiple rounds of primer synthesis occur at replication pause site [38]. Furthermore, these two ribonucleotides may not be processed by RNaseH and could be ligated with the properly processed end of the adjacent Okazaki fragment, thus catalyzing the imprint. Alternatively, incomplete processing of the RNA primers at the imprint site by RNaseH (which may be a result of replication pausing) may leave behind unprocessed 1-2 ribonucleotides to be ligated with the adjacent Okazaki fragments.

Interestingly, the residue D588 is conserved between S. pombe and metazonas, including human, mice, Xenopus [29], though the mammalian homologs have been shown to lack primase activity *in vitro* [31,39]. However, given the presence of an extra domain in the metazoan orthologs [39], a developmental control of primase activity through the extra domain may occur. Further studies are required to decipher such a possibility.

Furthermore, the reduced binding of Cdc23-M36 to Polα^+^ and still poorer binding to Polα^7-1^, may impede extension of RNA primers by the Polα^7-1^. This idea can explain the cumulative reduction we observed in imprinting/switching of *cdc23-M36, Polα^71^* double mutant on synthetic medium at 30°C. (The synthetic lethality observed on rich medium at 30°C may be due to requirement for some essential vegetative processes). The ability of Polα^D984N^ to restore imprinting in *swi7-1* mutant may be ascribed to possible restoration of recruitment of Polα^swi7-1^ to a Cdc23-bound template to help extend the RNA primers. The interaction of Cd23 with Swi1 also suggests that the unique primase activity may occur through interaction of Cdc23 with Polα and Swi1 at the pause site.

How imprinting is caused uniquely at *mat1* locus remains a puzzle. One possibility is that nucleotide sequence and unique DNA/nucleoprotein architecture at the *mat1* locus lead to a Swi1- and Swi3-dependent pause at the imprint site. This pause may facilitate localization of Cdc23 and Polα to persist at the imprint site long enough to synthesize a 2-ribonucleotide primer, which is then extended by the catalytic subunit of Polα during the elongation stage of DNA replication (Fig. 5). In addition, a weakened interaction of mutant Cdc23 with Polα protein might also cause reduced sequence specificity to install the imprint. Indeed, multiple break sites have been reported on both strands distal to the imprint site at the *mat1* locus in the *swi7-1* mutant [40]. It was proposed that Polα^swi7-1^ may not be able to guide the endonuclease to the correct site [40]. In light of the present results, we suggest that the mutant Polα^swi7-1^ may cause multiple ribonucleotide insertions near the imprint site, creating fragile sites, which lead to DNA cleavage during DNA preparation. Recently, lysine-specific demethylases Lsd1 and Lsd2 proteins have also been shown to control replication pause at *mat1* to facilitate efficient imprinting [41]. In sum our results suggest that Cdc23 plays a more direct role in imprinting than the previously described factors.

Strand-specific incorporation of ribonucleotides as a part of asymmetric DNA replication contributing to generation of developmental asymmetry of sister cells appears to be a unique process [4]. This apparent uniqueness may be because searching it requires knowing the locus that is differentially modified on one of the sister chromatids leading to a specific asymmetric cell division during development in metazoans. We recently reported that such a mechanism of asymmetric cell division indeed operates in evolutionarily distant yeast, *Schizosaccharomyces japonicus* [42].

Widespread incorporation of ribonucleotides has indeed been reported during mitochondrial DNA replication [43]. Recently, a low level of ribonucleotide incorporation by both Polα and Polδ has been demonstrated [44], raising the possibility of ribnucleotide insertion in DNA during differentiation or to facilitate recombination [45]. Similarly, fragile sites existing in mammalian genome occur predominantly in AT-rich sequences with a potential to form stem-loop structures, and are generated by the inhibitors of DNA synthesis that compromise replication by Polα [46]. Interestingly, Mcm10 plays an important role in mammalian development-its mutation causes defective embryonic cell proliferation and lethality in mice [47]. Interestingly, the primary defect occurs during morula to blastocyst stage when the inner cell mass (ICM) is formed, which ultimately produces different organs of the body. Thus, Mcm10 may play a role in cell differentiation during mammalian development. Thus, it would be interesting to investigate the role of primases in generation of DNA fragile sites, the incorporation of ribonucleotide insertions at these sites and their possible role in asymmetric cell differentiation during development in metazoans.

## Materials and Methods

### Strains and plasmids

The list of strains, plasmids and oligos used in this study are provided in Tables EV1, EV2 and EV3, respectively (Expanded View). Media and growth conditions employed were as described [49]. The per cent switching efficiency was determined by using the equation: 100 X [2(number of zygotes)/[2(umber of zygotes) + number of vegetative cells]. The transformant strains containing plasmid borne *Polα* or *cdc23* genes, expressed under the control of *nmt1* or *nmt41* promoters, were assayed for complementation on plates containing the repressor, thiamine. Under these conditions, leaky expression was observed from the *nmt1* promoter [50] (Ahmed and Singh, unpublished). For viability assays, the cultures of the required strains were normalized to the same OD_600_, serially diluted 10-fold and 5μl of each dilution was spotted on the required plates. Plates were incubated at 25°C for 5 days or 30°C for 3-4 days and then photographed.

### Southern hybridization

DNA was prepared by the normal method as described earlier [49]. DNA samples, isolated from yeast cells were digested *Hin*dIII endonuclease, were subjected to agarose gel electrophoresis, followed by Southern blotting and hybridization, as described earlier [11]. Alternatively, DNA was prepared from cells embedded in agarose plugs and subjected to restriction endonuclease digestion and alkali treatment as previously described [7]. The 10.6kb *mat1M*-containing *Hin*dIII fragment was used as a probe for Southern analysis. For experiments sown in Fig. EV2, donor deleted strain was used which only contains the *mat1* locus, while *mat2* and *mat3* loci have been deleted [51]. Normally the level of DSB is expected to be ~25of mat1 DNA signal when DNA is prepared by the conventional method. However, observe a higher level of ~35%, which may be ascribed to partial shearing of the mat1 DNA during preparation. Further, the level of DSB observed in alkali blots is around 10-12 % of *mat1* DNA. The 2-3 fold difference between the two methods of DNA preparation can be explained by the fact that the DSB fragments observed in blots of alkali-treated plugs detect only the imprinted strand due to single strand being cleaved while in case of DNA prepared by the normal method both strands are detected as the mat1 DNA is subjected to DSB.

### Iodine vapours-staining procedure of colonies used for quantifying switching efficiency

The homothallic cells, called *h^90^* cells, efficiently switch their mating type. Thereby, yeast colonies are composed of an equal proportion of P and M cells. Cells of opposite type mate under nitrogen starvation conditions, and the resulting “zygotic” diploid cell undergoes meiosis and sporulation to produce four haploid spores, called ascospores. The spores synthesize starch, but the vegetative growing cells do not. Because starch readily reacts with iodine vapours, when exposed to iodine vapours, efficiently switching colonies stain black in colour while those of reduced switching mutants stain lighter [49]. This procedure was used to test complementation of switching defective mutants.

### Primase Assay

Primase activity of Cdc23 was assayed according to et al. [48]. An oligo with the sequence: 5’..ACTTCGTCGACTTATAAAGACTGAAATGTAGCCTGAC….3’ was used for the assay.

## Acknowledgements

BS and KKB were supported by Senior Research Fellowships from Council of Scientific and Industrial Research, New Delhi, India. We are grateful to P. Nurse, J. Hurwitz, T. Wang and S. Kearsey for gifts of strains and plasmids. We thank L. Kaur for editorial help. This work received intramural support from Council of Scientific and Industrial Research, New Delhi, India. AK’s research is supported by the Intramural Research Program of The National Institutes of Health, Frederick National Laboratory for Cancer Research.

## Author Contributions

JS and AJSK conceptualized the research problem. JS, BS, KKB, ACK, and UU carried out the experiments. JSN and SS carried out the data analysis, JS and AJSK wrote and edited the manuscript.

## Conflict of Interest

The authors declare that there is no conflict of interest.

